# Signatures of omicron-like adaptation in early SARS-CoV-2 variants and chronic infection

**DOI:** 10.1101/2024.09.05.611549

**Authors:** Mark Tsz Kin Cheng, Mazharul Altaf, Jesu Castin, Ann-Kathrin Reuschl, Benjamin L. Sievers, Kimia Kamelian, Dejan Mesner, Rebecca B. Morse, Adam Abdullahi, Bo Meng, Kata Csiba, Cambridge NIHR Bioresource, Steven A. Kemp, Darren Martin, Clare Jolly, Christopher Ruis, Lipi Thukral, Ravindra K. Gupta

## Abstract

Omicron emergence represented a seismic event in the COVID-19 pandemic, demonstrating what was essentially antigenic shift in a virus that cannot reassort its genome as influenza can. Understanding the success of Omicron is essential, and yet we have little understanding of the biological underpinnings of its ability to accommodate diverse mutations and bring together deleterious mutations to generate a highly successful new serotype. Persistent SARS-CoV-2 infections are a likely source of new variants and may provide valuable insight into past and future evolutionary trajectories, in particular those involving allosteric interactions that defy genotype to phenotype prediction. Here we observe upper airway specific evolution of SARS-CoV-2 demonstrating fusion peptide (FP) domain mutation S:P812S adjacent to the S2’ cleavage site that emerged during a chronic infection in an immunocompromised individual. Indeed, this mutation had previously emerged in an ancestral B lineage as well as the delta variant lineage and transmitted successfully in populations globally, though remains uncharacterised. P812S in spike pseudotyped virus particles did not impact entry efficiency across cell lines expressing endogenous ACE2 and TMPRSS2. However, efficiency of spike cleavage at S1/S2 was reduced and molecular dynamics simulation demonstrated altered S1/S2 loop conformations that possibly impacted furin mediated cleavage. Consistent with impaired S1/S2 cleavage, and reminiscent of Omicron BA.1, cell-cell fusogenicity was severely impaired by introduction of P812S. The mutation also introduced significant perturbations to the FP region and also affected protomer-protomer packing. P812S conferred evasion of a neutralising monoclonal antibody targeting the fusion peptide, consistent with significant structural rearrangements in the FP region. Finally, P812S bearing viruses showed evasion of polyclonal neutralising antibodies in sera from vaccinated individuals at 32°C (simulating upper respiratory tract) and to a lesser extent at 37°C. Thus we report a novel mutational adaptation to the upper airway allowing enhanced immune evasion to fusion peptide targeting neutralising antibodies that also incurs a defect in ability to induce syncytia. These data shed light on the balance between upper airway adaptation/immune evasion by SARS-CoV-2, ability to induce syncytia formation, and disease severity given the established link between syncytia and severe COVID-19.

## Introduction

Omicron emergence represented a seismic event in the COVID-19 pandemic, demonstrating what was essentially antigenic shift in a virus that cannot reassort its genome as influenza can.

Understanding the success of Omicron is essential, and yet we have little understanding of the biological underpinnings of its ability to accommodate diverse mutations and bring together deleterious mutations to generate a highly successful new serotype. A key defining feature of Omicron in addition to immune evasion was the loss of ability of its spike to carry out cell-cell fusion.^1,2^ We do not have a clear understanding of which mutations or combinations are responsible given the multiple spike mutations observed across the protein. It is thought that this loss of syncytia formation was associated with the lower pathogenicity of Omicron, and that gradual reacquisition of spike mediated cell-cell fusogenicity by certain omicron subvariants correlates with increased severity.^3^

In virus producer cells, full length spike is cleaved by host furin to S1/S2 subunits which then reassociate through electrostatic interactions. The primary receptor for SARS-CoV-2 is ACE2 and following engagement with ACE2, the serine protease TMPRSS2 at the target cell membrane is able to cleave spike at the S2’ site, liberating the fusion peptide to initiate viral-cell fusion at the plasma membrane. The Omicron variant appears less dependent on TMPRSS2 and has a preference for endocytic entry where cathepsin carries out S2’ cleavage within endosomes.^2^ In addition we have observed a cleavage defect in virus producer cells and impaired cell-cell fusion.^2^ Given the broad expression of ACE2 across diverse tissues, it is unsurprising that infection of multiple organs occurs during COVID-19 infection. Local tissue environments likely exert specific selective pressure for replication. Viral divergence in different anatomical compartments has been reported, including respiratory compartments such as nasopharynx, sputum, trachea, lungs,^4^ and other compartments including plasma, heart, spleen, urine, and gut,^5,6^ consistent with differential selection pressure imposed by niches^12–15^. ACE2 and TMPRSS2 expression gradients exist between the upper-(URT) and lower-respiratory tract (LRT), and efficiency of TMPRSS2 use has varied with VOC. Omicron showed adaptation towards the URT^2,11^ and greater transmission potential. This tropism shift has been associated with reduced intrinsic disease severity.^12^ Affinity towards the URT also leads to differential exposure to environmental and immune selective pressures, creating a unique mutational spectrum which can be leveraged to determine the niche of pathogens.^5,11^ Another key difference is the difference in temperature, which has an impact on spike fusogenicity and thus entry efficiency.^13^

SARS-CoV-2 adaptation in persistent infection of immunocompromised hosts has been demonstrated in myriad studies.^14–17^ COVID-19 remains a particularly relevant threat for immunocompromised individuals, as COVID-19 vaccines have been shown to induce lower seroconversion rates,^18^ reduced neutralisation activity,^19^ and shorter durations of protection as compared to immune-competent individuals^20^. Here we leverage an intensively sampled case of chronic infection to report that a single mutation can have diverse phenotypic impacts and recapitulate some of Omicron’s key features. P812S in the fusion peptide region and also adjacent to the S2’ cleavage site was observed to arise in a sublineage of the Delta variant and in an early B lineage, yet remains uncharacterised^21^. We show that P812S causes changes in the fusion peptide (as one might expect), but also unexpected changes in the S1/S2 cleavage site that impairs cleavage. Moreover, the mutation severely impairs the ability of the spike to mediate cell-cell fusion and thereby we do not see syncytia formation. Syncytia formation is a hallmark of severe COVID-19.^22,23^ Importantly and unexpectedly, the mutation also confers a degree of escape from neutralising antibodies directed at the fusion peptide region. This is reflected in the reduced neutralisation of P812S bearing viruses by sera from vaccinated individuals, especially at low temperatures associated with the nasopharynx. Indeed, we first isolated the mutation on an early B lineage (pre-omicron virus) in a chronically infected individual from the upper airway and not in samples from the tracheal region-indicating this mutation may be an adaptation to the upper airway. These properties are reminiscent of Omicron and its adaptation to upper airway transmission with reduced severity correlating with impaired syncytia formation. Our work shows how careful genotypic and phenotypic study of chronic infections can shed light on the evolutionary biology of COVID-19 and.

## Results

### Clinical course of chronic infection

We previously reported an individual treated with rituximab for a lymphoma, who had a chronic COVID-19 infection with an early Wuhan-Hu-1+D614G SARS-CoV-2 variant (B.1.1.1) between April and August of 2020.^14^ The individual was treated with three courses of remdesivir / convalescent plasma and we previously documented immune escape conferred by D796H and viral infectivity enhancement conferred by delH69/V70. We obtained simultaneous samples from nose & throat swabs (NT) and lower endotracheal aspirates (ETA), though previously only data on throat swabs were analysed. ETA samples were only available starting from day 93 because the individual deteriorated and required intensive care admission and ventilatory support due to acute respiratory distress syndrome (ARDS)around day 90. From day 93 until day 101, five NT and four ETA samples were taken in total. The day 100 NT sample was removed from intra-host diversity analysis due to poor Illumina coverage and no available Ct value.

We first examined deep sequence data for the two anatomically proximate compartments of the nasopharynx and trachea. Using Shannon’s entropy as a measurement of viral diversity, we observed rises in diversity in NT samples when remdesivir or convalescent plasma were administered as the infection progressed, but no statistically significant difference in genetic diversity within the ETA compared to the NT samples **(Supplementary Figure 1a-d)**. Consistent with previous studies linking Ct values to viral diversity, the increase in viral diversity was accompanied by an increase in viral load.^5^ **(Supplementary Figure 1d)**

### Nucleotide spectral analysis reveals possible mixed upper and lower respiratory tract viral populations

The relative frequency of G>T mutations has previously been demonstrated to correlate with SARS-CoV-2 replication niche, with replication in the lung resulting in higher frequencies of G>T mutations.^11^ Omicron variants, which are more URT tropic than pre-Omicron variants,^2^ have lower G>T frequencies than pre-Omicron variants. **(Supplementary Figure 2a)** We therefore aimed to determine whether there was a difference in G>T frequencies between the NT and ETA isolates from the same patient. As expected given the short distance between these sites, we did not observe a difference between these niches **(Supplementary figure 2b)**; however, we noticed a clade of patient isolates that exhibited elevated numbers of G>T mutations **(Figure 1b,d)**. This clade contains both NT and ETA sequences from late in the course of infection (after day 93), suggesting spread of viruses within the patient across the respiratory tract. The elevation in G>T mutations amongst isolates in this clade suggests that this spread occurred from the LRT to the URT, consistent with deteriorating lung disease. This mixing of viral populations as the patient deteriorated clinically could have been related to the physical conduit provided by the endotracheal tube or air movement associated with mechanical ventilation.

**Figure 1:**
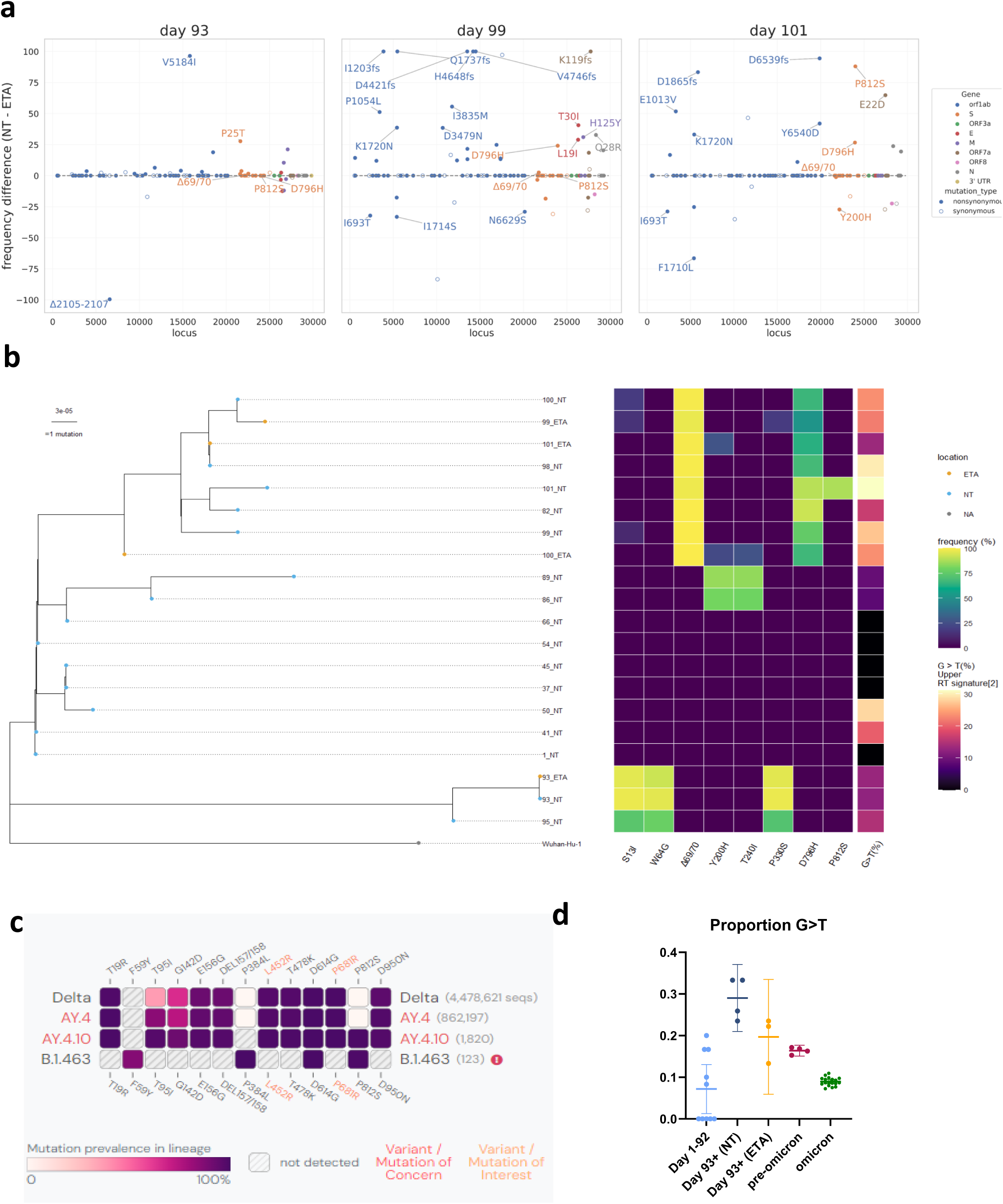
Genetic changes over time and across niches during chronic COVID-19. a) The absolute difference in percentage prevalence of mutations in patient X1 plotted against the SARS-CoV-2 genomic map. Nonsynonymous mutations (filled) with >20% difference are annotated. P812S is found on day 101 NT compartment. b) Maximum-likelihood phylogenetic tree of patient X1 with the day of sampling and location (blue=nose and throat swabs, orange= endotracheal aspirate) indicated. Middle panel shows the Spike mutation frequency foundin each sample, and right panel the proportion of G to T mutation in each sample. c) Global prevalence data taken from Outbreak.info on 7^th^ April 2024. left: Global prevalence of S:P812S per PANGO lineage. Right: Spike mutations which occurs at greater than 75% in one of Delta (B.1.617.2),AY.4, AY.4.10, B.1463. d) Proportion of G to T mutations (a lower airway mutation signature) in patient between day 1 to 92, day 93 onwardsfrom NT samples, day 93 onwards from ETA samples, and pre-omicron and omicron variants. Day 93 and Day 95 were excluded as they are separated from the main clade. The data for omicron and pre-omicron variants are aggregated data from available sequences of the UShER SARS-CoV-2 phylogenetic tree. Data are plotted with mean and 95% CI.

### Multiple nonsynonymous spike and non-spike mutations arise following therapies for COVID-19

We further examined the changes in mutation frequencies between the NT samples compared to the ETA samples. Although the long branch with S:S13I, S:W64G, and S:P330S emerged, there is limited divergence in variant frequencies between NT and ETA samples on day 93 immediately after remdesivir administration and two units of convalescent plasma infusion. On day 95, divergent mutations arose for Spike (Δ69/70, D796H, P812S) and other genes, including ORF1ab (E1013V), Envelope (T19I), Nucleocapsid (Q28R), and Membrane (H125Y) **(Figure 1a, Supplementary figure 1e, Supplementary figure 2c)**. This is likely due to selection pressure imposed by the convalescent plasma and/or remdesivir treatment.^17^ Of note, the spike mutation **P812S** emerged on day 101 at a frequency of 88.0% on day 101 in the NT sample, but not in the ETA sample. **P812S**was accompanied by a 98.1% frequency of Δ69/70 and a 89.7% frequency of 796H (Figure 1a, **Supplementary figure 1d,3**), suggesting linkage of these three mutations on the same viral genomes.

At the time of this study (7^th^ April 2024), P812S had a low prevalence in the Global Initiative on Sharing All Influenza Data (GISAID) database (<0.5%, n =12380) **(Table 1, supplementary figure 3)**. P812S was most abundantly found in Delta variant viruses (n = 5744), but was also found in the Alpha variant (n = 1819) and the major Omicron lineages BA.1 (n=610), BA.2 (n=602), BA.4/5 (n=1181), and XBB (n=507) viruses **(Table 1, Supplementary Figure 3)**. Though P812S is a minority in these major lineages, over 99.6% of AY.4.10 sublineage sequences (1801 out of 1808) and 99.2% of B.1.463 sublineage sequences (122 out of 123) contained the P812S mutation **(Figure 1c)**, indicating it was a defining mutation in these lineages and arguing for successful transmission of P812S-bearing viruses.^24–26^ Indeed, from the start of the pandemic to August 2023, spike codon 812 had an average dN/dS ratio of 2.47 (*p*=0.184), suggesting positive selection (i.e., selection favouring changes away from the P amino acid state). (https://observablehq.com/@spond/sars-cov-2-global-genomic-selection-2019-aug-2023). Statistical evidence (p<0.01 with the IFEL method) of positive selection at codon 812 was most marked between May to November 2021 when the Delta variant was most prevalent, corroborating the beneficial effect of diverging on a Delta genomic background from P812 to alternative amino acids such as P812S under at least some epidemiological conditions. **(Supplementary figure 4).**

**Table 1:** Lineage of P812S-containing sequences on GISAID.

| **scorpio_call** | **Aggregate Group** | **frequency** | **percentage** |
| --- | --- | --- | --- |
| Alpha (B.1.1.7-like) | Alpha (B.1.1.7-like) | 1818 | 14.68% |
| B.1.1.7-like+E484K | Alpha (B.1.1.7-like) | 1 | 0.01% |
| Delta (AY.4-like) | Delta (AY.4-like) | 2903 | 23.45% |
| Delta (AY.4.2-like) | Delta (AY.4-like) | 21 | 0.17% |
| Delta (B.1.617.2-like) | Delta (B.1.617.2-like) | 2820 | 22.78% |
| Omicron (BA.1-like) | Omicron (BA.1-like) | 610 | 4.93% |
| Omicron (BA.2-like) | Omicron (BA.2-like) | 602 | 4.86% |
| Omicron (BA.5-like) | Omicron (BA.4/5-like) | 1127 | 9.10% |
| Omicron (BA.4-like) | Omicron (BA.4/5-like) | 54 | 0.44% |
| Omicron (XBB.1-like) | Omicron (XBB.x-like) | 210 | 1.70% |
| Omicron (XBB.1.5-like) | Omicron (XBB.x-like) | 162 | 1.31% |
| Omicron (XBB-like) | Omicron (XBB.x-like) | 109 | 0.88% |
| Omicron (XBB.1.16-like) | Omicron (XBB.x-like) | 26 | 0.21% |
| Gamma (P.1-like) | Other | 167 | 1.35% |
| A.23.1-like | Other | 61 | 0.49% |
| Iota (B.1.526-like) | Other | 20 | 0.16% |
| Epsilon (B.1.429-like) | Other | 18 | 0.15% |
| Beta (B.1.351-like) | Other | 16 | 0.13% |
| Epsilon (B.1.427-like) | Other | 7 | 0.06% |
| Mu (B.1.621-like) | Other | 7 | 0.06% |
| Lambda (C.37-like) | Other | 6 | 0.05% |
| B.1.617.1-like | Other | 4 | 0.03% |
| Eta (B.1.525-like) | Other | 1 | 0.01% |
| Theta (P.3-like) | Other | 1 | 0.01% |
| Zeta (P.2-like) | Other | 1 | 0.01% |
| Unassigned | Unassigned | 1590 | 12.84% |
| Probable Omicron<br>(Unassigned) | Unassigned | 10 | 0.08% |
| Omicron (Unassigned) | Unassigned | 8 | 0.06% |
|  | Total | 12380 |  |

All SARS-CoV-2 isolates bearing P812S as of 7<sup>th</sup> April 2024, aggregated by Scorpio assigned lineage.

We stratified by variant and proportionally subsampled 247 SARS-CoV-2 sequences containing P812S and inferred a maximum-likelihood tree **(Supplementary Figure 3; GISAID EPI_SET_240114wm)**. We annotated the sequences based on the accompanying mutations and noticed that, apart from the D614G mutation that arose early in the pandemic, there were no mutations consistently found to occur with P812S.

As P812S was found only in the NT sample, we hypothesised that P812S conferred an advantage in that niche. The proximity to the S2’ cleavage site and fusion peptide, in addition to the exposed position of P812 when Spike is in the open position **(PDB: 6XM4, Figure 2a)** prompted us to study the infectivity, cleavage, cell-cell fusion and immune evasion properties of P812S mutants.

**Figure 2:**
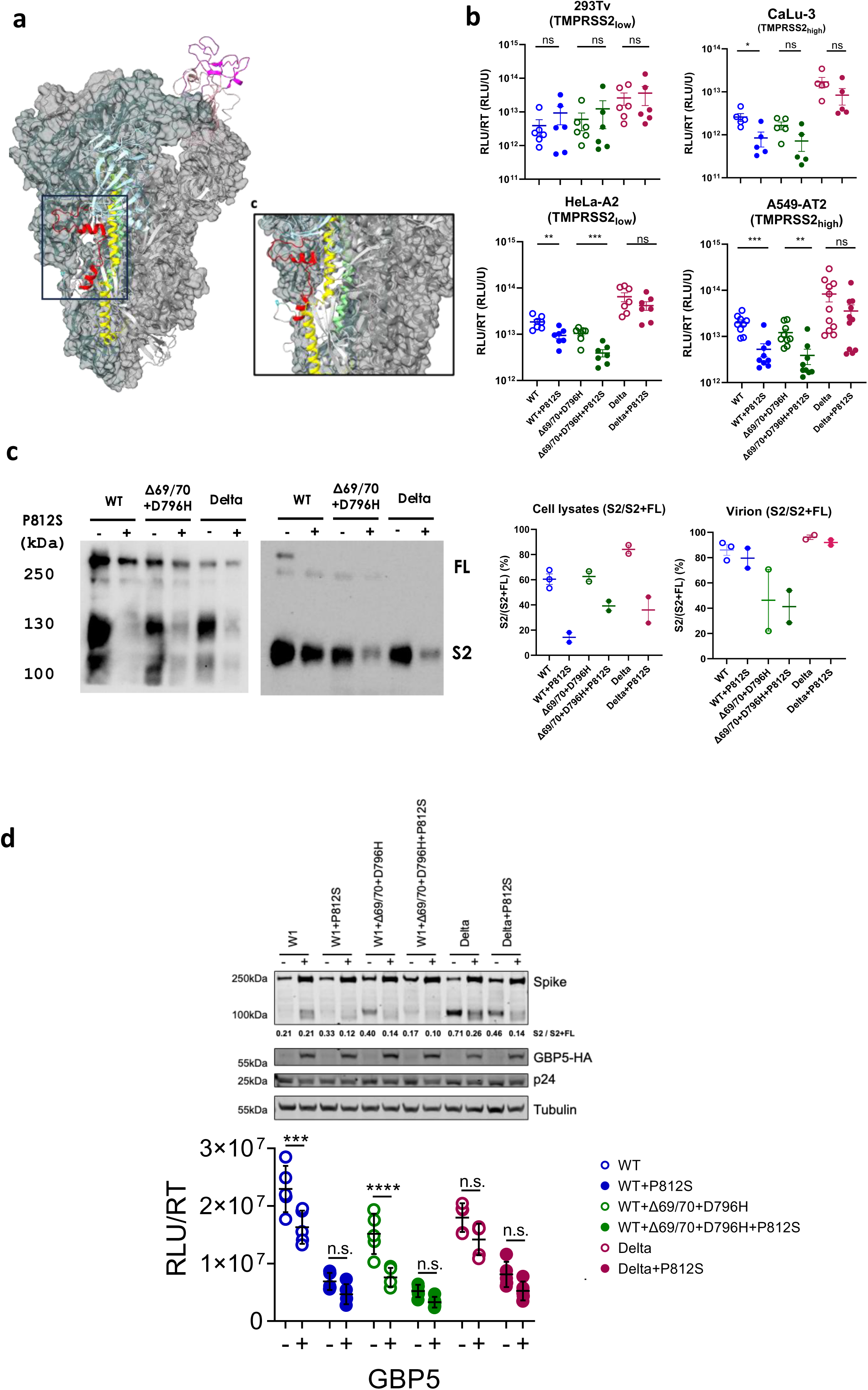
P812S bearing spikes maintain infectivity in cell lines expressing endogenous levels of ACE2/TMPRSS2 but show impaired S1S2 cleavage and reduced sensitivity to GBP5. **a**) Structure of SARS-CoV-2 Spike (PDB: 6XM4) where two subunits are in the “RBD down” conformation (gray surfaces) and one subunit is in the “RBD up” conformation (coloured ribbon). Domains are coloured as follows: NTD (light blue), RBD (pink), RBM (magenta), P812 (cyan), Fusion peptide (red), HR1 (yellow), HR2 (green). b) PV entry of WT (Wuhan-Hu-1+D614G), WT+P812S, WT+Δ69/70+796H, WT+Δ69/70+796H+P812S, Delta, Delta+P812S introduced to ACE2-high, TMPRSS2-low 293Tv-A2; Hela-A2; and ACE2-high, TMPRSS2-high A549-AT2, Calu-3 lung cells at 37°C. Data are supplemented by mean ± standard error of n = 3–8 technical replicates. Statistical analysis was performed using unpaired student’s t-test. c) Western blot analysis of spike cleavage in producer HEK293Tv cell lysate and isolated pseudovirus. S2 to FL spike ratio were analysed by densitometry (ImageJ), and S2/S2+FL cleavage proportion was calculated for virions and cell lysates. d) GBP sensitivity of Spike-Gag-PVs (produced in the presence or absence of GBP5 on ACE2/TMPRSS2-A549. Top panel shows western blot of transfected producer cell lysates probed with antibodies to S2, HA, p24 and tubulin. Bottiom panel shows PV infectivity (RLU/ RT U) and is representative of two independent experiments with 5-6 measurements each Statistical analysis was performed using one way ANOVAwith

### P812S leads to a general decrease in S1/S2 cleavage and variable cell entry efficiency

As P812S is directly upstream (N-terminal) to the S2’ cleavage site (R815/S816) and the S2 viral fusion peptide (FP, 816-855),^27–29^ we hypothesised that P812S may directly impact spike glycoprotein S2 cleavage and viral entry. To test this, we introduced the P812S mutation into Wuhan-Hu-1 + D614G (WT), WT with Δ69/70 + 796H mutations, as well as Delta B.1.617.2. We further incorporated Delta and Delta+P812S in the panel because P812S was predominantly found in Delta and previous genomic-epidemiological studies showed P812S^21^ further increases the likelihood of Delta breakthrough infections^30,31^. We then performed infectivity assays using pseudoviruses with and without the P812S mutations on cell lines that varied in ACE2 and TMPRSS2 expression levels. We observed little impact of P812S in cells expressing endogenous levels of receptors **(Figure 2b)**.

However, P812S was associated with a significant deficit in cell entry efficiency in cell lines where ACE2 or TMPRSS2 were overexpressed **(Figure 2b, Supplementary Figure 5).** P812S-associated entry defects persisted across 32°C, 37°C, and 39°C, a temperature which approximates pyrexia**. (Supplementary Figure 6)**.

As the cleavage of S1/S2 has been shown to correlate with infectivity/entry/transmission efficiency, to explore this further, we performed western blot for full length (FL) and S2 spike on 293T producer cell lysate and purified virions.^32^ We noticed that P812S was associated with suboptimal S1/S2 cleavage in both producer cell lysate and virion, demonstrated by decreased ratio of S2 to FL spike (**Figure 2c**). The P812S-induced reduction of S2 in cell lysate and released virions suggests that P812S may play a role in stabilising the S protein and slowing S1 shedding and it appears that cleaved spike is preferentially incorporated into viral particles as we observed previously.^32^ A reduction in S1/S2 cleavage is reminiscent of the change observed from Delta to Omicron variant^2^, which also showed tropism shift away from cells expressing TMPRSS2 and impaired cell-cell fusion in Omicron.^11,33^ We next sought to confirm whether P812S confers an entry route preference shift, as seen in the transition of global sequences from the TMPRSS2-reliant plasma membrane entry favoured by the Delta variant to cathepsin-reliant endosomal entry favoured by the Omicron variant (**Table 2**). P812S was associated with a <2-fold resistance to both E64d (cathepsin inhibitor, inhibiting endosomal entry) and camostat (serine protease inhibitor, including TMPRSS2, inhibiting plasma membrane entry). In summary we did not observe substantial shift in efficiency of receptor or coreceptor usage for P812S.

**Table 2.** half maximal effective concentration (EC50) of camostat and E64d.

| Spike presented on pseudotyped Virus | EC50 (µM) |  |
| --- | --- | --- |
|  | camostat | E64d |
| WT | 16.85 | 1.99 |
| WT+P812S | 23.58 | 3.24 |
| Δ69/70+D796H | 17.69 | 1.79 |
| Δ69/70+D796H+P812S | 23.76 | 4.50 |
| Delta | 19.45 | 2.11 |
| Delta + P812S | 23.59 | 3.87 |

GBP5 is an interferon inducible protein that inhibits furin cleavage and was shown to impact HIV Env cleavage by furin^34,35^ as well as SARS-CoV-2.^36^ Given we observed an S1/S2 cleavage defect for P812S bearing spike, we wished to explore what impact GBP5 might have on furin cleavage of the P812S spike. To test this, we co-expressed HA tagged GBP5 along with spikes bearing a range of mutations and packaging/reporter genome plasmids. We isolated PV and infected target A549-A2T2 lung cells lines. We observed that GBP5 reduced infectivity of each virus tested, and that the effect was smaller for PV bearing P812S, consistent with cleavage that was already impaired at S1/2. Western blot analysis reflected these findings (**Figure 2d**).

### Significant conformational rearrangements in the fusion peptide and S1/S2 cleavage site regions associated with P812S

To probe structural changes in P812S Mutant in SARS-CoV-2 spike (S) protein, we generated two molecular dynamics simulations of glycosylated wildtype and P812S spike variant based on experimentally resolved cryo-EM structure 7A94 with 1RBD up bound to ACE2 (**Supplementary Figure 6**). The global arrangement of the SARS-CoV-2 spike (S) protein, which is structurally stabilized through the packing interactions of its three protomers, was significantly perturbed upon the introduction of the mutation. The protomer area formed by three CH domains increased from 86.8 A^2^ to 91.8 A^2^ upon the introduction of P812S mutation, leading to altered spatial organization of the protomers (**Supplementary Figure 6b**). Comparative structural analysis revealed increased dynamic fluctuations, with pronounced flexibility observed in the mutant, primarily originating from the NTD, FP and S1-S2 regions (**Supplementary Figure 6c**). Notably, the fusion peptide (FP) region of the SARS-CoV-2 spike (S) protein, a short, hydrophobic segment within the S2 subunit, is positioned just downstream of the S2’ cleavage site (R815) and spans residues 816–833. To gain deeper insights into the structural changes surrounding this region, we extended our analysis to encompass residues 788–855, allowing us to capture broader conformational dynamics that may impact fusion efficiency and viral entry. Furthermore, the region of S1/S2 proteolytic cleavage site (residues 669–696) also showed greater structural compaction in the P812S mutant compared to the WT, suggesting potential alterations in cleavage accessibility and spike activation dynamics. To understand the residues that are responsible for the local perturbation at the FP region (**Supplementary Figure 6**), we monitored the backbone conformational changes and computed rotamer shift with respect to WT (explained in methods). The shift is majorly elevated in the loop flanking FP helix, which is comprising of residues 830 to 855, as shown in Supplementary Figure 6e-f. These structural modelling studies provide insight into our observations of impaired S1/S2 cleavage, consistent with our understanding that proteolytic cleavage at the S2’ site may be impacted by P812S mutation.

### P812S confers evasion of serum neutralising antibodies

Multiple broadly neutralising antibodies have been identified to interact with amino acids on the N-terminal side of the FP (816-655), including P812.^37–39^ In addition, P812 has been identified as a highly exposed epitope.^40^ Therefore, we hypothesised that P812S may confer a degree of immune escape due to selective pressure imposed by convalescent plasma treatment. We tested the neutralisation sensitivity of Wuhan-Hu-1 + D614G (WT), WT+P812S, WT+Δ69/70+D796H, and WT+ Δ69/70+D796H+P812S using a previously reported pseudovirus (PV) system.^41^. We tested 20 stored vaccinee sera (average age 66.0, 55% M) one month after second-dose AstraZeneca ChAdOx1 COVID-19 vaccine.^42,43^

We confirmed previous observations^14^ showing modest immune escape by the double mutant Δ69/70+D796H, with a 1.2-fold decrease in geometric mean serum neutralising titres between WT and WT+Δ69/70+D796H **(Figure 3)**. P812S on a WT+Δ69/70+D796H backbone was associated with a further loss of sensitivity to neutralising serum antibodies **(Figure 3)**. This suggests that the immune evasion capabilities conferred by D796H are enhanced by P812S.

**Figure 3:**
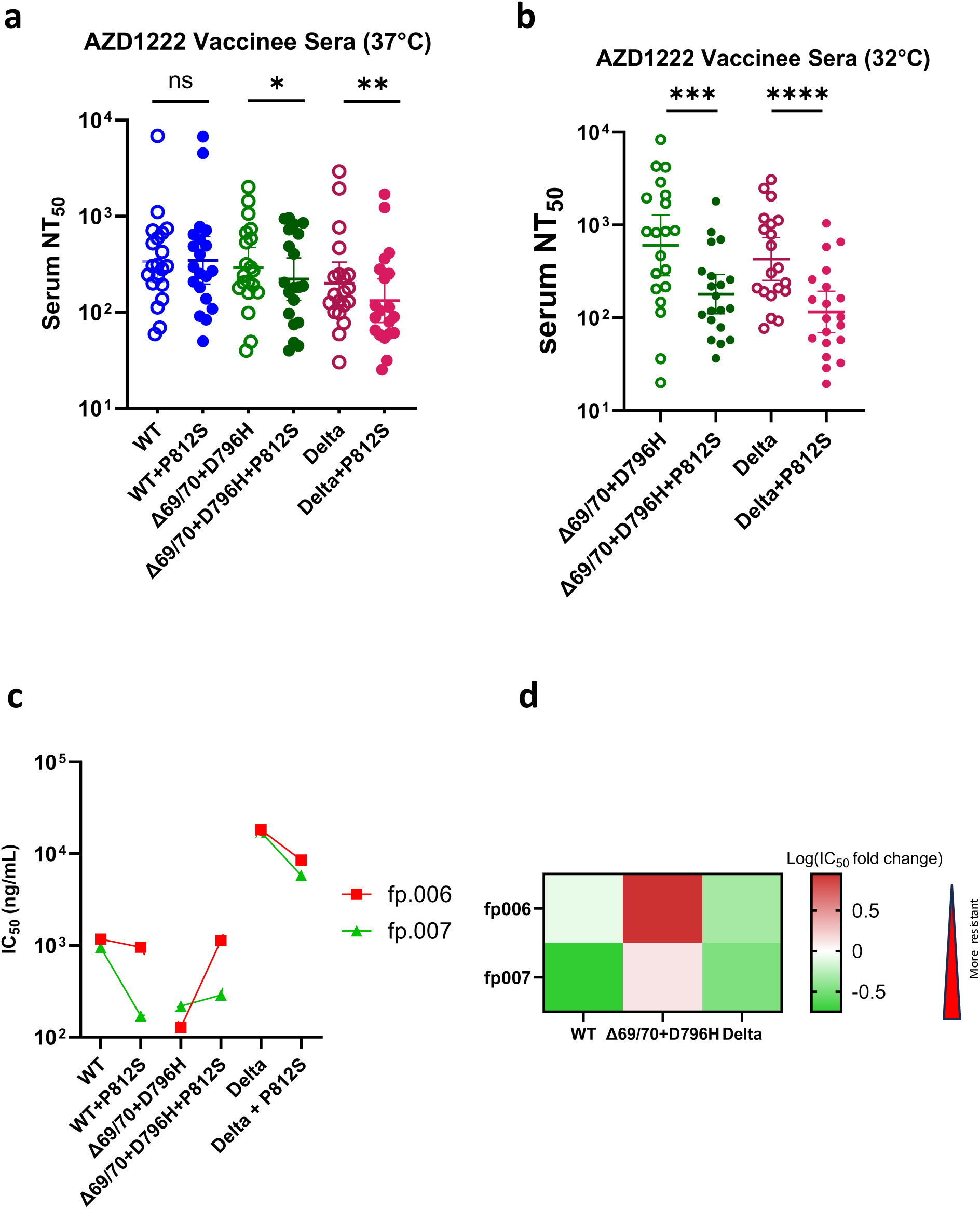
P812S confers escape from polyclonal sera from vaccinated individuals and neutralising antibodies against the fusion peptide. a, b) Neutralization by second dose adenoviral vector (AZD1222) vaccinee-elicited sera against of WT (Wuhan-Hu-1+D614G), WT+P812S, WT+Δ69/70+796H, WT+Δ69/70+796H+P812S, Delta, Delta+P812S pseudotyped virus. Pseudovirus was co-incubated with sera for one hour at 37°C (a) or 32°C (b). Reciprocal geometric mean titre shown with 95% confidence interval. Statistical analysis was performed using ratio paired t test corrected for multiple comparison using the Holm-Šídák method.; *P <.05, **P <.001, ***P <.0001. **c**) IC50 of fp.006 (red), and fp.007 (green) incubated with pseudovirus pseudoviruses corresponding to the indicated mutations. Results are representative of 2 technical replicates. d) The log-transformed fold change induced by adding the P812S to each spike backbone is represented in the heatmap.

To further understand the impact of the P812S mutation, we produced Delta pseudovirions with and without the P812S mutation. Using sera one month after two doses of AstraZeneca ChAdOx1-S, both baseline serum neutralising titres against Delta and Delta+P812S were lower than WT (Table 3).

**Table 3.**
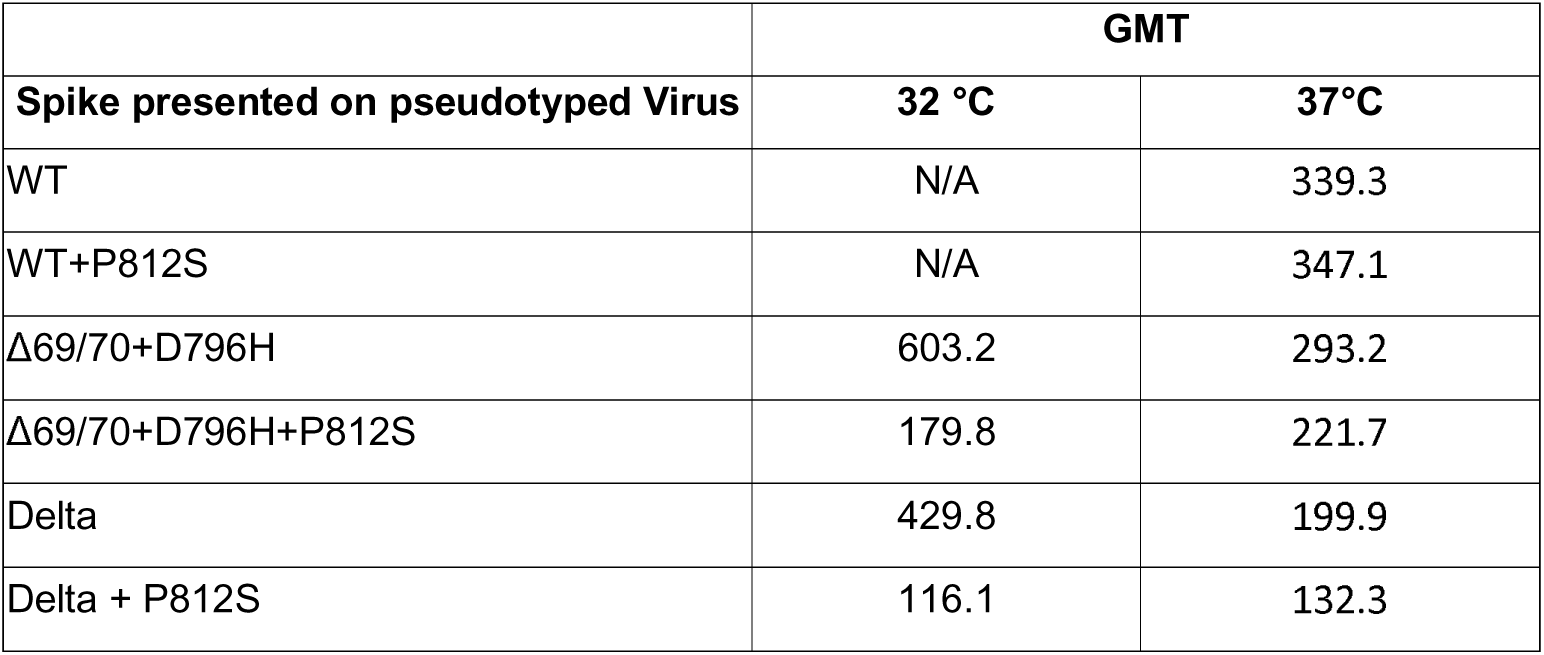
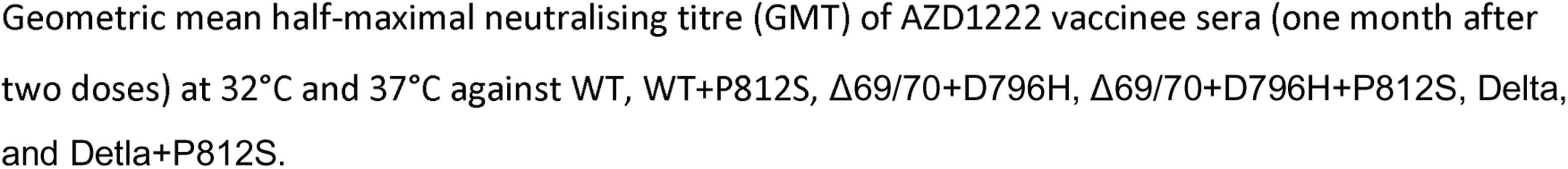
Geometric mean half-maximal neutralising titre (GMT) of AZD1222 one month post second dose vaccinee sera of P812S mutants.

| Spike presented on pseudotyped Virus | GMT |  |
| --- | --- | --- |
|  | 32 °C | 37°C |
| WT | N/A | 339.3 |
| WT+P812S | N/A | 347.1 |
| Δ69/70+D796H | 603.2 | 293.2 |
| Δ69/70+D796H+P812S | 179.8 | 221.7 |
| Delta | 429.8 | 199.9 |
| Delta + P812S | 116.1 | 132.3 |

Geometric mean half-maximal neutralising titre (GMT) of AZD1222 vaccinee sera (one month after two doses) at 32°C and 37°C against WT, WT+P812S, Δ69/70+D796H, Δ69/70+D796H+P812S, Delta, and Delta+P812S.

There was a significant decrease in serum neutralisation GMT of the Delta P812S mutant compared to Delta, consistent with immune evasion **(Figure 3)**. In summary, at 37°C, in both the Delta and WT+Δ69/70+D796H backbones, P812S incorporation conferred a degree of resistance to serum neutralisation. To better simulate the upper airway environment, we tested PV expressing WT+Δ69/70+D796H or Delta +/-P812S against sera/monoclonal antibody at 32°C.^44^ The serum geometric mean titre for 32°C increased for all viruses by between x1.5 to x2.0 fold compared to 37°C. (Table 3) At 32°C, P812S also conferred some resistance to vaccinee sera by x1.4 fold in the background of WT+Δ69/70+D796H and x1.3 fold in the Delta background, respectively **(Figure 3).**

Finally, we wished to explore the impact of P812S on susceptibility to Sotrovimab treatment, given it is was the only licensed therapeutic monoclonal during the chronic infection case. Sotrovimab (derived from S-309) (GSK) is a monoclonal class 3 antibody which targets a highly conserved epitope within the Spike receptor binding domain (RBD).^45^ In agreement with previous literature, Sotrovimab demonstrated similar neutralisation titres with serum against WT and Delta, ^46^ and P812S had limited effect on sotrovimab susceptibility, with IC50 fold differences less than two fold in the presence versus absence of P812S **(Table 4, Supplementary Figure 7)**.

**Table 4.** half-maximal inhibitory concentration (IC50) of SARS-CoV-2 Spike targeting monoclonal antibodies.

| Spike presented on<br>pseudotyped Virus | IC50 (ng/mL) |  |  |  |
| --- | --- | --- | --- | --- |
|  | Sotrovimab (32°C) | Sotrovimab (37°C) | fp.006 | fp.007 |
| WT | NA | 46.8 | 1173 | 953 |
| WT+P812S | NA | 31.03 | 960 | 170.8 |
| Δ69/70+D796H | 37.30 | 45.38 | 127.8 | 218.3 |
| Δ69/70+D796H+P812S | 14.79 | 55.04 | 1135 | 288.4 |
| Delta | 8.60 | 45.09 | 18286 | 17195 |
| Delta + P812S | 9.71 | 79.56 | 8545 | 5818 |

IC50 of Sotrovimab fp.006, fp.007 against WT, WT+P812S, Δ69/70+D796H, Δ69/70+D796H+P812S, Delta, and Delta+P812S.

### P812S confers evasion of neutralising antibodies targeting the fusion peptide

To gain some understanding of the mechanism of P812S on immune evasion, two monoclonal antibodies were tested. We used fusion-peptide targeting recombinant monoclonal antibodies *fp.006* and *fp.007* from Bianchini and colleagues.^29^ In brief, *fp.006* recognises a conserved fusion peptide epitope that is only exposed upon ACE2 binding. IC50 reduction conferred by P812S for WT (x1.22) and Delta (x2.14) is modest for the ACE2-binding dependent fp.006, compared to the IC50 increase for Δ69/70+D796H (x8.88). This suggests that in on the background of Δ69/70+D796H the P812S decreased sensitivity of the FP epitope to neutralising antibodies in sera, but not on a WT Wu-1 background.

Antibody *fp.007* binding affinity and neutralising capacity is independent of ACE2 binding. *fp.007* demonstrated a significant 5.58-and 2.96-fold decrease in IC50 upon P812S introduction on both WT and Delta backbones, respectively, with little difference upon P812S introduction to the Δ69/70+D796H double mutant. **(Figure 3**, **Table 3)**. Taken together it appears that FP antibodies are likely to have driven selection of P812S especially as P812S induced significant conformational changes in the FP region.^47^ In order to probe this further, we performed MD simulations of the Δ69/70+D796H double mutant with and without P812S (**Figure 4a**). The average inter-protomer distance increased to 15.2 Å in the presence of P812S along with background mutations (**Figure 4b**). Although the perturbations were not as high as observed in P812S without background mutations (**Supplementary Figure 6**), we captured elevated fluctuations at the FP and S1/S2 regions. We computed ΔRMSF (RMSF_WT_-RMSF_MUT_) to assess the changes in the fluctuation patterns upon mutation. In the FP region, the P812S, along with background mutations, introduces local instability with up to 1 Å increase in RMSF (**Figure 4c-d**). We could capture relatively increased fluctuations in the S1/S2 region (up to ∼3Å) in P812S + background mutations. While the presence of P812S imparted instability to FP and S1/S2 sites, the lack of P812S (Δ69/70+D796H double mutant) showed no major changes in the fluctuations (**Figure 4e-g**). These observations support the observed escape from FP targeting antibody and the reduced S1/S2 cleavage seen for the triple mutant.

**Figure 4.**
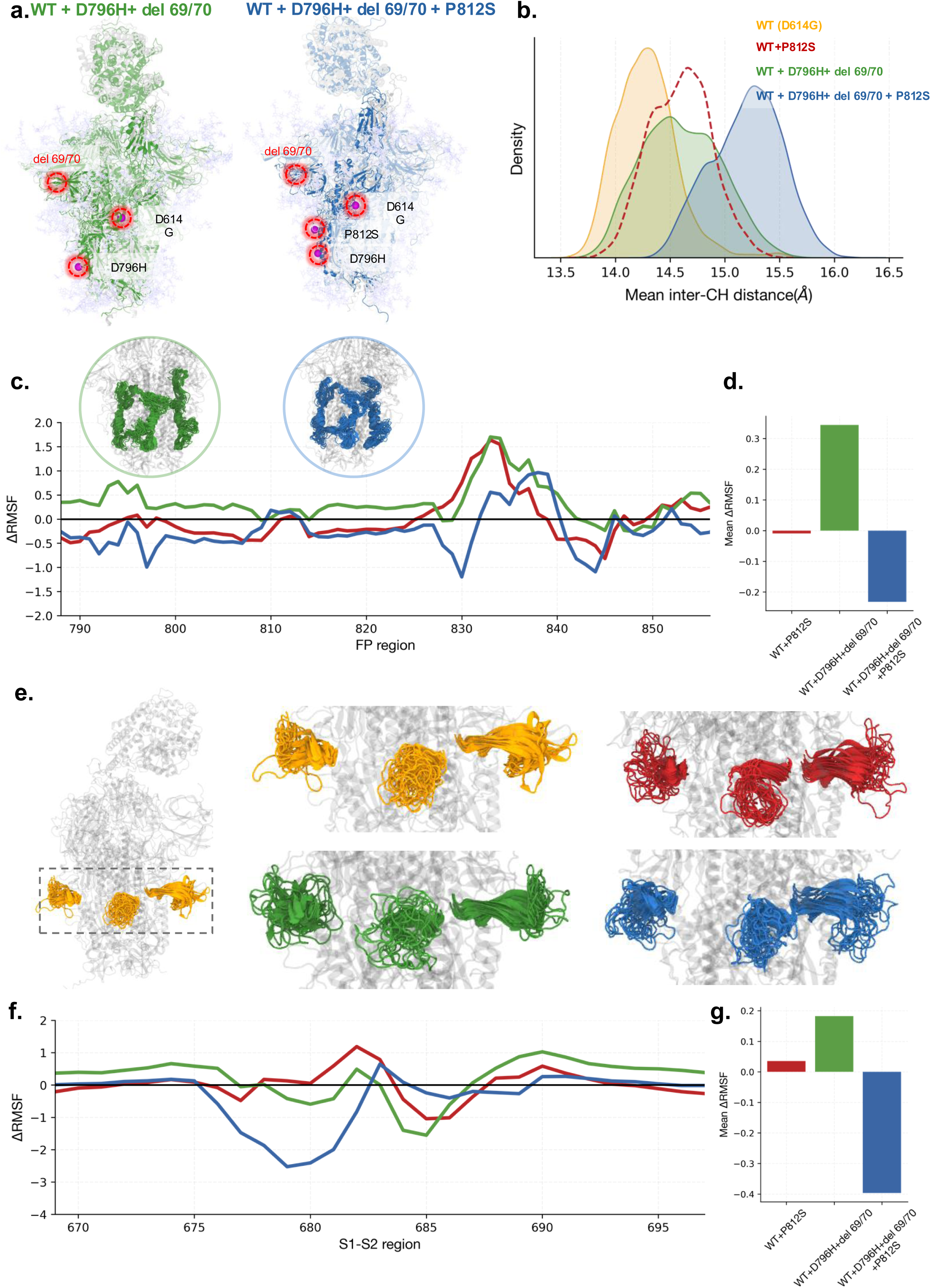
Presence of P812S imparts instability to the S1/S2 cleavage and fusion peptide regions. a) The snapshots highlight the background double mutant (D796H+ del 69/70) with (right) and without (left) P812S. The position of mutations is highlighted. b) The average inter-CH distance distributions are plotted for WT and all three mutants. c) ΔRMSF (RMSF_WT_-RMSF_MUT_) is plotted for residues of the FP region. The snapshots highlight the conformational ensemble of FP region seen in the background mutations with and without P812S substitution. d) The average ΔRMSF from the FP region is plotted for all the mutants. e) The snapshots highlight the dynamics of S1/S2 region in WT and three mutations. f) ΔRMSF is plotted for S1/S2 region highlighting all three mutants. g) The average ΔRMSF underlying S1/S2 residues is plotted for all the mutants.

### P812S severely impairs cell-cell fusion

The ability to induce syncytia via cell-cell fusion is a known property of type I virus glycoproteins, including HIV-1 Env, RSV F and SARS CoV-2 S proteins.^48^ This process requires processing of S1/S2 in the producer cell and activity of a serine protease in the target cell to cleave S2’ and release the fusion peptide. Syncytia are thought to be a correlate of disease severity in COVID-19.^49^ We previously showed that Omicron BA.1 shifted biologically from Delta, demonstrating severely reduced cell-cell fusion activity in vitro, and that this appeared to correlate with S1/S2 cleavage.

Subsequent Omicron variants such as BA.4 have regained some fusogenicity.^50^ Given the impact of P812S on both S1/2 cleavage and on FP structural dynamics, we measured the impact of P812S on cell-cell fusion activity using a split GFP system. As expected, Delta spike fusion was highest (**Figure 5**), followed by the double mutant D796H and del69/70 and then WT. Surprisingly, we observed a severe defect induced by the P812S mutation on each background.

**Figure 5:**
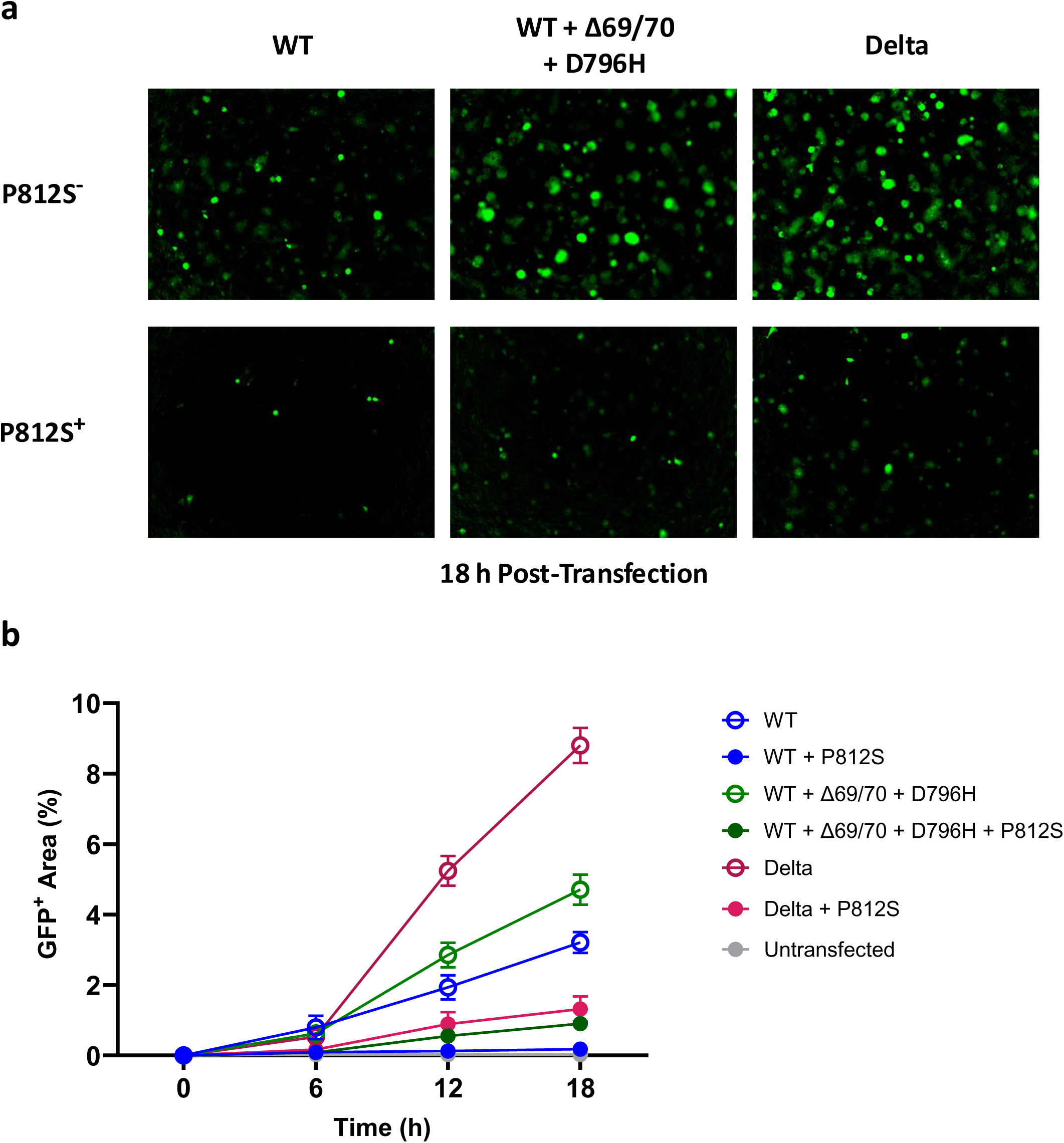
Cell-cell fusion kinetics of P812S bearing SARS-CoV-2 spike proteins. Vero-GFP1-10 co-incubated with 293T-GFP11 cells transfected with DNA plasmids encoding SARS-CoV-2 spike proteins bearing a range of mutations. a) Representative images of spike-mediated cell-cell fusion in WT (Wuhan-Hu-1+D614G), WT+P812S, WT+Δ69/70+796H, WT+Δ69/70+796H+P812S, Delta, Delta+P812S at 18 hours post-transfection. Images were captured using the IncuCyte SX5 Live-Cell Analysis System (Sartorius). b) Quantification of spike-mediated cell-cell fusion showing the percentage of GFP+ area to the total cell area at 0-, 6-, 12-, and 18-hours post-transfection. Cell-cell fusion was determined as the proportion of green area to the total phase area and was calculated with the IncuCyte analysis software (Sartorius). Data are mean ± s.e.m. from four fields of views at each time point and are representative of two independent experiments.

## Discussion

Persistent SARS-CoV-2 infections are a likely source of new variants and may provide valuable insight into past and future evolutionary trajectories, in particular those involving allosteric interactions that defy genotype to phenotype prediction. Here we observe upper airway specific evolution of SARS-CoV-2 demonstrating fusion peptide (FP) domain mutation S:P812S adjacent to the S2’ cleavage site that emerged during a chronic infection in an immunocompromised individual. This mutation had also previously emerged in an ancestral B lineage as well as the delta variant lineage and transmitted successfully. P812S in spike pseudotyped virus particles did not impact entry efficiency across cell lines expressing endogenous ACE2 and TMPRSS2. However, efficiency of spike cleavage at S1/S2 was reduced and molecular dynamics simulation demonstrated altered S1/S2 loop conformations that likely impacted furin mediated cleavage. We noted that GBP5, an interferon inducible restriction factor that inhibits furin mediated cleavage, had a smaller impact on inhibition of P812S bearing pseudotyped viruses, consistent with a pre-existing S1/S2 cleavage defect in P812S bearing spikes.

Consistent with this observation and reminiscent of Omicron BA.1, cell-cell fusogenicity was severely impaired by introduction of P812S. The mutation also introduced significant perturbations to the FP region and protomer-protomer packing. On the other hand, P812S conferred evasion of a neutralising monoclonal antibody targeting the fusion peptide, consistent with significant structural rearrangements in the FP region. As the S2 subunit and FP region are strong candidates for pan-sarbecoviruses broadly neutralising antibody (bnAb) therapy and vaccines,^29,37,38,51^ our work shines light on potential functional deficits associated with immune escape mutations.

Finally, P812S bearing viruses showed evasion of polyclonal neutralising antibodies in sera from vaccinated individuals at 32°C (simulating upper respiratory tract) and to a lesser extent at 37°C. We propose that the selection of P812S in the upper airway compartment is an immune evasion adaptation that came at the cost of cell-cell fusion and entry efficiency in some cells overexpressing ACE2/TMPRSS2, possibly mediated through impaired S1/2 cleavage in P812S-bearing spikes.

Interestingly, this URT adaptation occurred on the background of within-host evolution suggestive of adaptation to the LRT based on mutational spectra analysis. Thus, it is plausible that the LRT population migrated to the URT given the intense inflammation in the lower airway in the present case. Although we were unable to separate URT and LRT isolates in mutational spectra due to the similarity of sequences at the consensus level from our patient, comparison of patient sequences and global sequences of pre-Omicron and Omicron lineages is suggestive of progression from lower to upper airway. Our work demonstrates the importance of studying compartmentalisation of viral populations, especially in persistent infection. Monitoring these mutations that have epistatic effects could potentially assist in prediction of mutations in future circulating strains. The driving force for mutations in the immunocompromised host is significantly different and allows for unique combinations, especially given suboptimal antibody-driven selection pressure.

It is interesting and notable that a single mutation can have multiple unexpected phenotypic impacts. These observations provide clues as to the events leading to the emergence of the highly successful Omicron variant, highlighting the role of allosteric impacts of immune escape mutations – for example on cleavage sites - and the importance of mutational context. Indeed, the dogma that viruses become less pathogenic over time is widely quoted, and yet the evidence for this is weak. If syncytia formation indeed is a correlate of disease severity, our data provide one route as to how such a process might occur for SARS-CoV-2, although ultimately the ‘holy grail’ is a step by step understanding of how the Omicron variant selected individual mutations.

## Limitations of the study

Our study was based on lentiviruses pseudotyped with different SARS-CoV-2 spikes. Although consistency has been observed between pseudotyped virus system and the live-virus system for the study of virus entry,^42,52^ there may still be potential effectors only present in the live-virus system.^2^ Confirmation of our finding in a live-virus system would be desirable.

## Acknowledgement

This work was funded by a Wellcome Senior Fellowship to RKG (WT108082AIA). This research was supported by the NIHR Cambridge Biomedical Research Centre (NIHR203312). A-KR and CJ are funded by a Wellcome Investigator award to CJ (223065). LT is thankful to the funding support from the DBT/Wellcome Trust India Alliance (IA/21/2/505925). The views expressed are those of the authors and not necessarily those of the NIHR or the Department of Health and Social Care. We thank the valuable discussion with Dr Maximillian Muenchhoff on SARS-CoV-2 HLA and CD8 T cell epitope evasion analysis. We also thank Dr Filippo Bianchini, Ms Jasmine Chantal Cantergiani, and Dr Davide F. Robianni for sending samples and expression plasmids of fp.006 and fp.007 antibodies.

## Author Contributions

Conceptualization, M.T.K.C. and R.K.G.;

Methodology, M.T.K.C., D.M.,C.R.,B.M. M.A;

Investigation, M.T.K.C., A.M., D.M., C.R., B.L.S., K.K., R.B.M., M.A., K.C. and A-K.R.

Writing – Original Draft, M.T.K.C.;

Writing – Review & Editing, M.T.K.C, M.A, D.M., C.R., R.K.G. A-K.R. and C.J.

Funding Acquisition, R.K.G;

Resources, R.K.G;

Supervision, B.M., S.A.K., R.K.G., C.J.

## Methods

### Clinical-sample collection and sequencing

Sequencing of SARS-CoV-2 viral DNA extracted from serial samples from the LRT (sputum and endotracheal aspirate), and URT (throat and nasal swab) was described previously in Kemp et al.^14^ In short, nucleic acid extraction was done from 500 μL sample using the easyMAG platform (Biomerieux). Sequencing amplicons were prepared with the ARTICnetwork v.3 protocol and diluted to 2 ng μl^−1^ and 25 μl (50 ng) were used as input for each library preparation reaction, before sequencing using the Illumina MiSeq platform.

Sera from recovered patients in the COVIDx study were used for testing of neutralization activity by SARS-CoV-2 mutants.

### Sequence data processing and analysis

Illumina sequencing data from Kemp et al. was re-analysed.^14^ Bam files were downloaded from Kemp et al. (BioProject <u>PRJNA682013</u>), mapped to Wuhan-Hu-1 (NC_045512.2) by minimap v2.28 (arguments-ax sr),^53^ sorted and indexed by samtools v1.20,^54^ and variants were called by BCFtools v1.18 functions mpileup (minimum base quality q30), and call (arguments--multiallelic-caller--variants-only).^55^ Mutations identified by BCFtools were filtered by a minimum depth cutoff of 20x before annotations by snpEff v5.2 using the publicly available NC_045512.2 database.^56^

### Genetic diversity and viral load analysis

The genetic diversity of each sample was estimated using Shannon Entropy based on the frequency of each iSNV, assuming that all iSNVs are independent from each other.

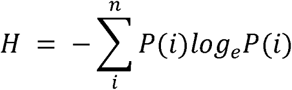

Where *P(i)* is the iSNV percentage frequency at gene loci *i*.

Shannon Entropy of NT samples collected prior to day 93, NT samples collected on day 93 and after, and ETA samples collected on day 93 and after were compared using student’s t-test. For Ct values, Welch’s t-test was performed.

### Phylogenetic analysis and lineage assignment

Excluding duplicates, and low coverage sequences, we searched GISAID for all complete coverage SARS-CoV-2 sequences with the Spike_P812S substitution on the 7^th^ April 2024. All sequence encoding Spike-S812 were aligned to MN908947.3 by mafft (v7.525,--6merpair--keeplength--addfragments), and then were assigned lineages using Pangolin (V4.3).^57^ The major lineages were grouped by Scorpio (V0.3.19). We stratified the sequences based on their major lineage designations and then performed stratified random sampling with proportional allocation to major lineages to obtain a subset of 250 P812S-containing sequences using the R package Rsample v1.2.1.

We constructed a maximum-likelihood tree from patient X1 and the 250 sequence sample using IQ-Tree v2.2.5 employing the GTR+F+I model of nucleotide substitution (model with the lowest Bayesian Information Criterion identified by IQ-Tree ModelFinder) with clustering support assessed by 1000 rounds of ultrafast bootstrapping.^58^

### Signals of natural selection in the SARS-CoV-2 S-gene

Mutational sites found in the patient, the Alpha variant, and the Delta variant, together with sites of neutralisation escape mutations were chosen for natural selection analysis using the internal-branch fixed effects (IEFL) method implemented in HYPHY.^59^ All globally sampled near-full length SARS-CoV-2 sequences are sampled between December 2019 and March 2022 were binned in three-monthly intervals (labelled according to the last month of sampling within a bin) and analysed separately within the context of the entire Spike open reading frame: an analysis setup needed for both phylogenetic context and for informing the nucleotide and codon substitution models upon which the IEFL method relies. Only codon substitutions mapped to internal phylogenetic tree branches were considered in these analyses since mutations mapping to terminal branches would have likely experienced insufficient durations of selection to yield meaningful evolutionary information. The approximate dominant-variant era during the COVID-19 pandemic where indicated based on results from Amodio and colleagues.^60^

### Structure Analysis

Coordinates for the SARS-CoV-2 Spike trimer complex with two RBD down and one RBD up was from PDB: 6XM4-Chain A to C, and domains definition from multiple references were used.^29,61–63^ Graphics describing the structures were made in UCSF Chimera X V1.8.^64^

### Mutational Spectrum analysis

We calculated the mutational spectrum of mutations within each patient sample using the mutation files generated above. The korimuto pipeline within MutTui v2.0.2^65^ was run on each sample independently using NC_045512.2 as the reference genome. The proportion of G>T mutations within each sample was calculated as the number of G>T mutations divided by the total number of mutations.

### Generation of P812S Spike mutants

Amino acid substitutions were introduced into a codon-optimised pCDNA3.1_SARS-CoV-2 WT, Δ69/70, Delta **(B.1.617.2, EPI_ISL_2378732)**, and Omicron (**EPI_ISL_7418017**) Spike Plasmid as previously described,^2,14,31^ using the QuikChange Lightning Site-Directed mutagenesis kit, as per manufacturer’s protocol (Agilent Technologies).

### Molecular dynamics simulations

The spike protein in ‘1-RBD-up’ state in complex with ACE-2 was considered for molecular dynamics simulation. The D614G spike was considered as the wild type (WT) protein. Along with D614 spike, three different mutant systems were prepared – i) WT+ P812S, ii) WT+D796H+del 69/70 and iii) WT+D796H+del 69/70+P812S. The spike protein was glycosylated as given in Casalino et al., 2020. The systems were solvated with TIP3P water and further neutralized with the counter ions. All the simulations were carried out with Gromacs version 2021.3 incorporating the CHARMM36 all-atom force field (Abraham et al., 2015). A grid spacing of 0.16 nm was used along with the fourth-order cubic interpolation for Particle Mesh Ewald (PME) summation. The van der Waals cut-off was set to 1.2 nm. The systems were initially subjected to energy minimization with steepest descent method. Temperature and pressure were maintained at 310 K and 1 bar using V-rescale thermostat and Parrinello-Rahman barostat, respectively. The numerical integration of equation of motion was computed every 2 fs and the coordinates of the system were saved 100 ps. Each system was simulated for 1 μs. The structures were visualized and rendered with PyMOL (Schrödinger, LLC, 2015). All basic distance and RMSD/RMSF calculations were performed in Python with MDAnalysis library.

### Computing backbone rotamer shift with respect to WT

To track the residues that has changed their backbone dihedral conformations, we computed a comparative metric called rotamer shift. We discretized the observed backbone dihedral angle into *cis* and *trans* states as given in Singh et al., 2017. We quantify the rotamer shift for the residue R_i_ with the following equation.

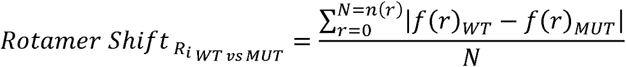

The r denotes the rotamer state of the dihedral. For backbone dihedrals, the rotamer states include Cis and Trans. Thus, N is the total number of rotamers for a dihedral. Each of the rotamer fractions f(r) (fraction of a rotamer r of a residue throughout the trajectory) is computed for WT and the mutant and subtracted. The rotamer shift is then defined as the average of absolute value of these differences in the rotamer fractions. The resulting shift varies from 0 (no changes in rotamer fractions) to 1 (complete change in rotamer fractions). We computed rotamer shift in *ψ* and *ϕ* angles for each residue. The maximum of two was finally considered to represent the dihedral change occurred at a residue. The dihedral angles were calculated with the gromacs command *gmx rama*. The rotamer assignment and the shift calculations were performed with the in-house Python script.

### Pseudovirus preparation

Pseudovirus were prepared by transfecting HEK293Tv cells with 1µg of p8.91 HIV-1 gag-pol expression vector, 1.5 µg of pCSFLW, and 1µg of plasmid expressing the respective spike protein using a 3:1 ratio of Fugene HD transfection agent (Promega).^41,66^ After 48 hours of incubation at 10% CO_2_, 37°C, viral supernatant was collected and filtered through a.45 µm filter and stored at-80°C until further use. The 50% tissue-culture infectious dose (TCID_50_) of SARS-CoV-2 pseudovirus was determined using the Steady-Glo Luciferase assay system (Promega).

For GBP5 sensitivity assays, pseudoviruses were prepared as described previously.^36^ In brief, HEK293T cells were transfected the indicated spike plasmids, p.8.91 and pCSLW in the presence of 40ng GBP5 vector^34,35^ or a corresponding empty vector control using Fugene6 (Promega). After 48-72h, culture supernatants were collected and 0.45 µm filtered.

### Standardization of Virus Input by reverse transcriptase activity using SYBR Green-Based Product Enhanced RT-PCR Assay (SG-PERT)

The reverse transcriptase activity of pseudovirus preparations was determined by quantitative polymerase chain reaction (qPCR) using a SYBR green-based product-enhanced PCR assay (SG-PERT) as previously described.^67^ Briefly, undiluted supernatants were lysed in a 1:1 ratio in a 2x lysis solution (40% glycerol v/v, 0.25% Trition X-100 v/v, 50 mM KCl, Ribolock RNase inhibitor 0.8 U/mL, TrisHCl 100mM, buffered to pH 7.4) for 10 minutes at room temperature.

### Western blot analysis

For virus pellets, the filtered viral supernatant was centrifuged at 15,000 rpm for 120 min to pellet virions. For cell lysates, transfected HEK293Tv cells were resuspended in PBS, before centrifugation at 1500 rpm for 5 minutes. The cell pellet was then lysed in 400 µL cell lysis buffer (Cell Signaling Technology Cat#9803) and sonicated for 1 minute.

All samples are further lysed and reduced in Laemmli reducing buffer (1M Tris-HCl (pH6.8), SDS, 100% glycerol, β-mercaptoethanol and bromophenol blue). This was followed by electrophoresis on SDS 4-12% Bis-Tris protein gels (Thermo Fisher Scientific) under reducing conditions. Proteins on the gel are transferred by electroblotting onto polyvinylidene difluoride (PVDF) membranes. The SARS-CoV-2 spike proteins and pseudotype virus capsid were visualized using a ChemiDoc MP imaging system (Biorad) or Odyssey Cxl Infrared Imager (Licor) using a rabbit anti-Spike polyclonal (Thermo Fisher Scientific, Cat# PA1-41165, RRID:AB_1087210, LOT:WE3286564, 1 in 4000), mouse anti-p24 monoclonal (BEI Resources; NIAID Cat# ARP-3537, RRID:AB_3086785, 1 in 3000), mouse anti-Spike monoclonal (GeneTex 1A9, 1:1000), mouse anti-HA-tag (Biolegend, 16B12 1:2000), rabbit anti-p24 (CFAR ARP432, 1:2000) and mouse anti-alpha-tubulin (Sigma DM1A, 1:1000).

Chemiluminescence was detected using Bio-Rad Clarity Western ECL Substrate (Cat#1705061), or Cytiva Amersham ECL Prime (Cat#RPN2232), and the band intensity is quantified using Fiji Software (ImageJ v.153k).

### EC50 of entry inhibitors in A549-ACE2-TMPRSS2 cells with PV

A549-ACE2/TMPRSS2 (A549-A2T2) cells were treated with either E64d (Tocris) or camostat (Sigma-Aldrich) for 2 h at each drug concentration before the addition of a comparable amount of input pseudotyped viruses (approximately 500,000 RLU). The cells were then left for 48 h before addition of lysis buffer + luciferase substrate (Promega) and read on a Glomax plate reader (Promega).

Normalisation was performed using an inhouse script, and EC_50_ values were calculated using a nonlinear regression model of inhibitor dilution vs. normalized response with a variable slope in GraphPad Prism 10.

### Vaccinee serum and monoclonal antibody pseudotype virus (PV) neutralization assay

Previous work have shown that IC_50_ obtained from spike pseudotype assays quantitatively correlate with neutralisation activity measured using an authentic SARS-CoV-2 neutralization assay.^52^

Virus neutralisation assays were conducted on HeLa cells stably expressing ACE2 (HeLa-A2) using SARS-CoV-2 spike pseudotyped virus expressing firefly luciferase as previously described.^14^

Sera were first heat-inactivated at 55°C for an hour. Serial dilutions of sera or purified monoclonal antibodies are then incubated with pseudotyped virus for 1h at 37°C. Freshly trypsinized and quenched HEK293T-A2T2 or HeLa-A2 cells were subsequently added to each well. After 48h incubation at 37°C and 5% CO_2_, luminescence was measured using a Bright-Glo luciferase assay system (Promega) and neutralisation was calculated relative to cell-only and cell-and-virus-only controls. Normalisation was performed using an inhouse script, and IC_50_ values were calculated using a nonlinear regression model of inhibitor dilution vs. normalized response with a variable slope in GraphPad Prism 10.

---

**Supplementary Figure 1.**
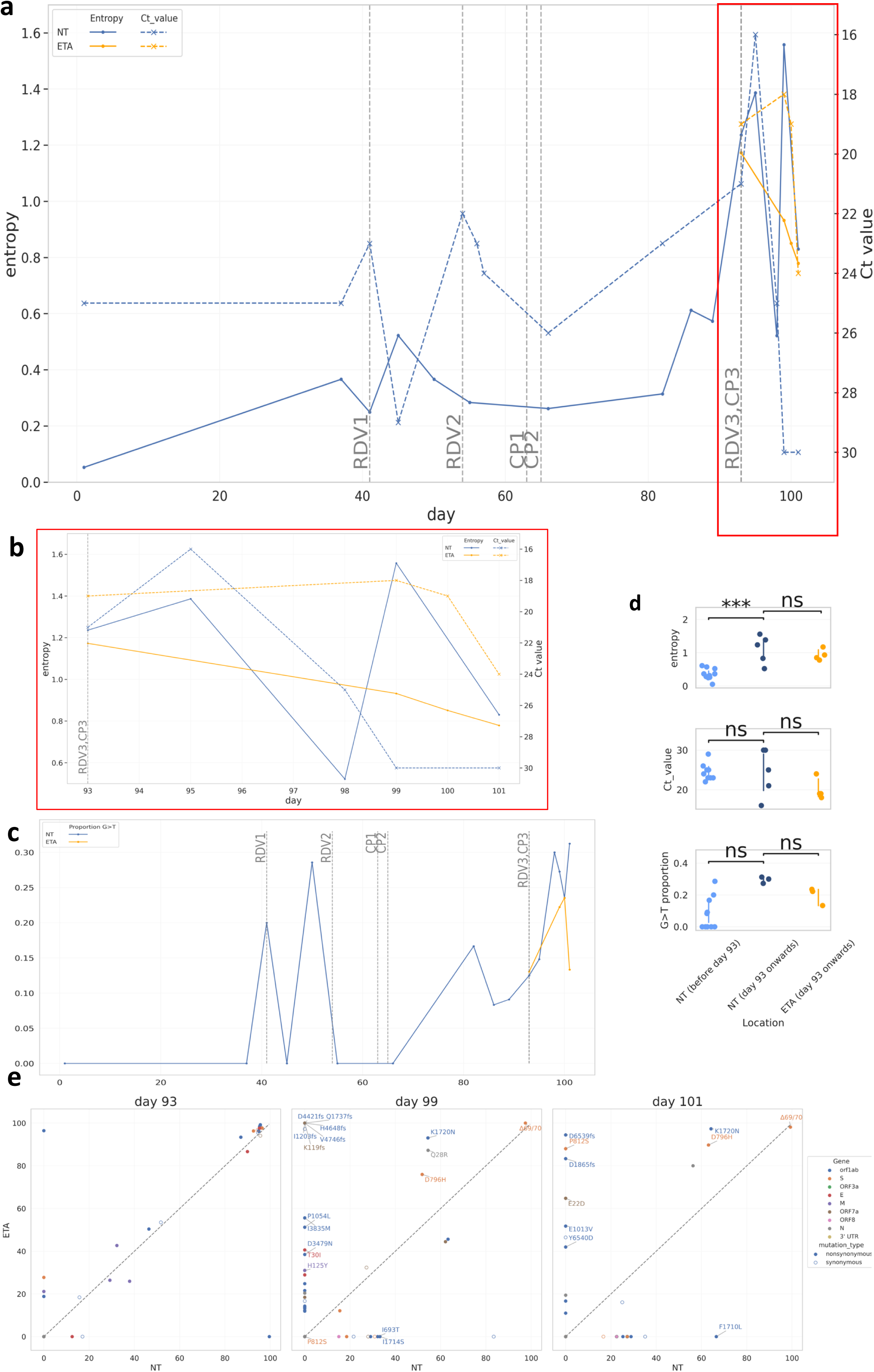
: Viral load and Intra-host evolution dynamics. a) Shannon’s entropy (solid line, left y-axis) and Ct value (dotted line, right y-axis) of nose and throat (N+T, blue) and endotracheal aspirate (ETA, yellow) samples, zoomed in and rescaled from day 93 to day 101 (b). CP, convalescent plasma; RDV, Remdesivir. c) proportion of G to T mutation from day 1 to day 101. Day 93 and Day 95 were excluded as they are separated from the main clade. d) Dot plot (with 95% confidence interval) of Shannon’s entropy (top) and Ct value (bottom) for NT samples from day 1-92, NT samples from day 93-101, and ETA samples from day 93 to 101. Student’s t-test is performed for entropy whilst Welch correction was performed for Ct value in lieu of unequal variance. e) The percentage prevalence in N+T plotted against ETA. Nonsynonymous mutations (filled) with >30% difference are annotated. The grey reference line represents equivalent prevalence between NT and ETA.

**Supplementary Figure 2:**
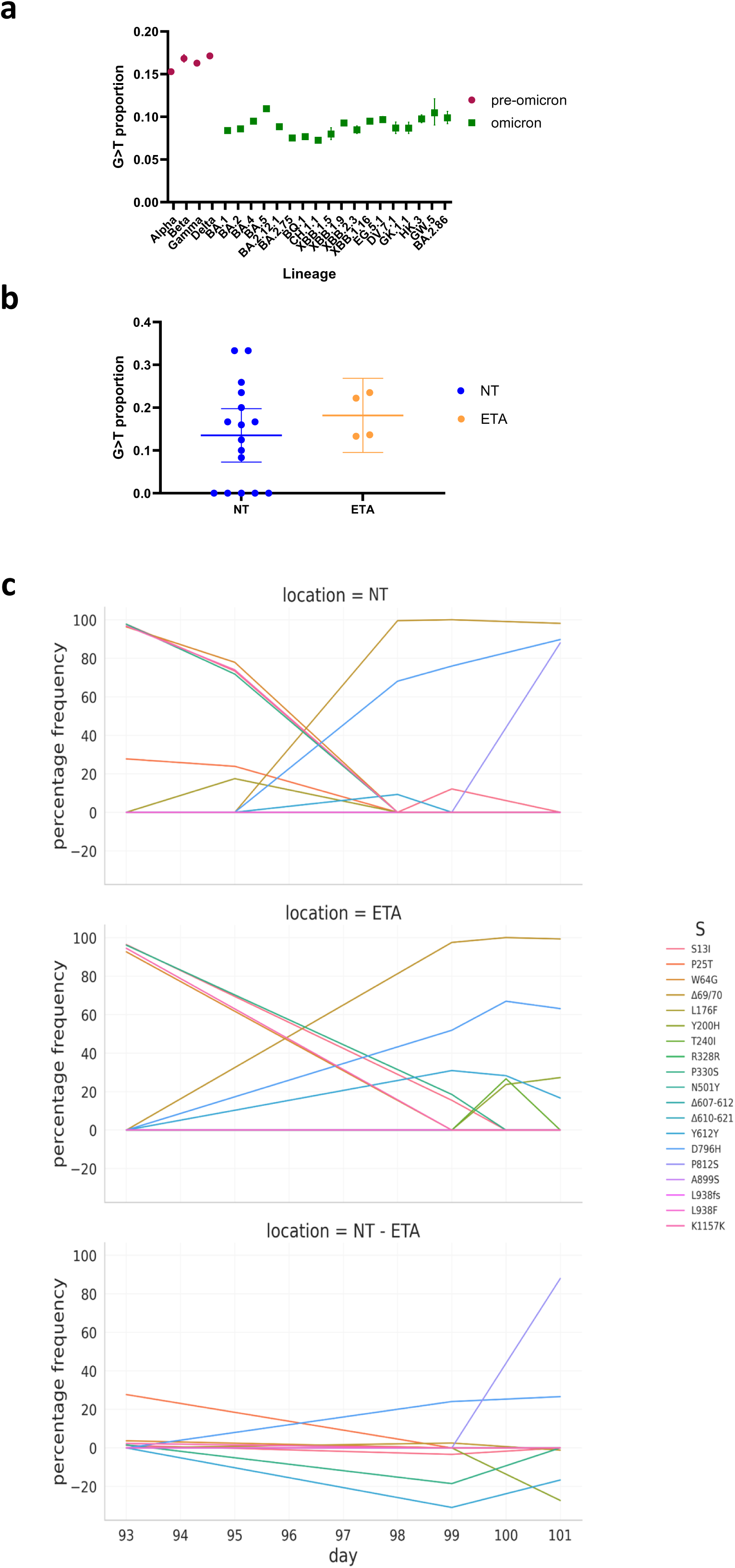
Breakdown of mutational spectra and viral iSNV dynamics in the nasal and endotracheal compartments. The G to T proportion of a) globally circulating lineages in chronological order and b) of patient sample broken down by NT and ETA samples across all timepoints. C) Spike mutation frequency in N+T (top), ETA (middle), and absolute percentage difference (bottom) between day 93 and 101.

**Supplementary Figure 3.**
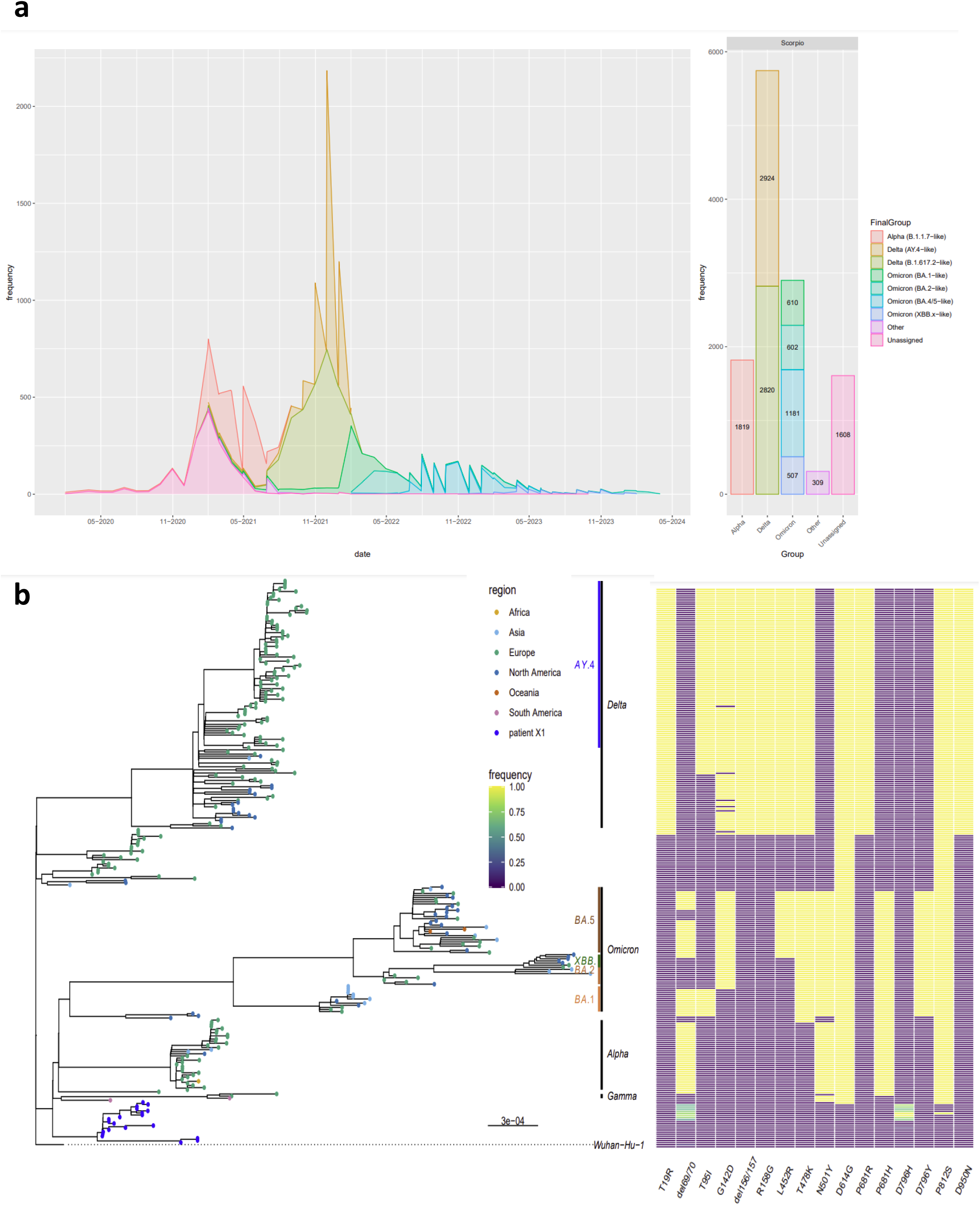
: Global Context of P812S containing SARS-CoV-2 global variants. a) The global prevalence of SARS-CoV-2 sequences with the S:P812S mutation in different variants defined by Scorpio as of 7^th^April 2024. b) Left panel - Maximum Likelihood tree (model: GTR+F+I) of patient X1 and 247 variant-stratified subsampled sequences. Tips are labelled by region. Right panel – Aligned heatmap of Spike mutations of all sequences.

**Supplementary Figure 4:**
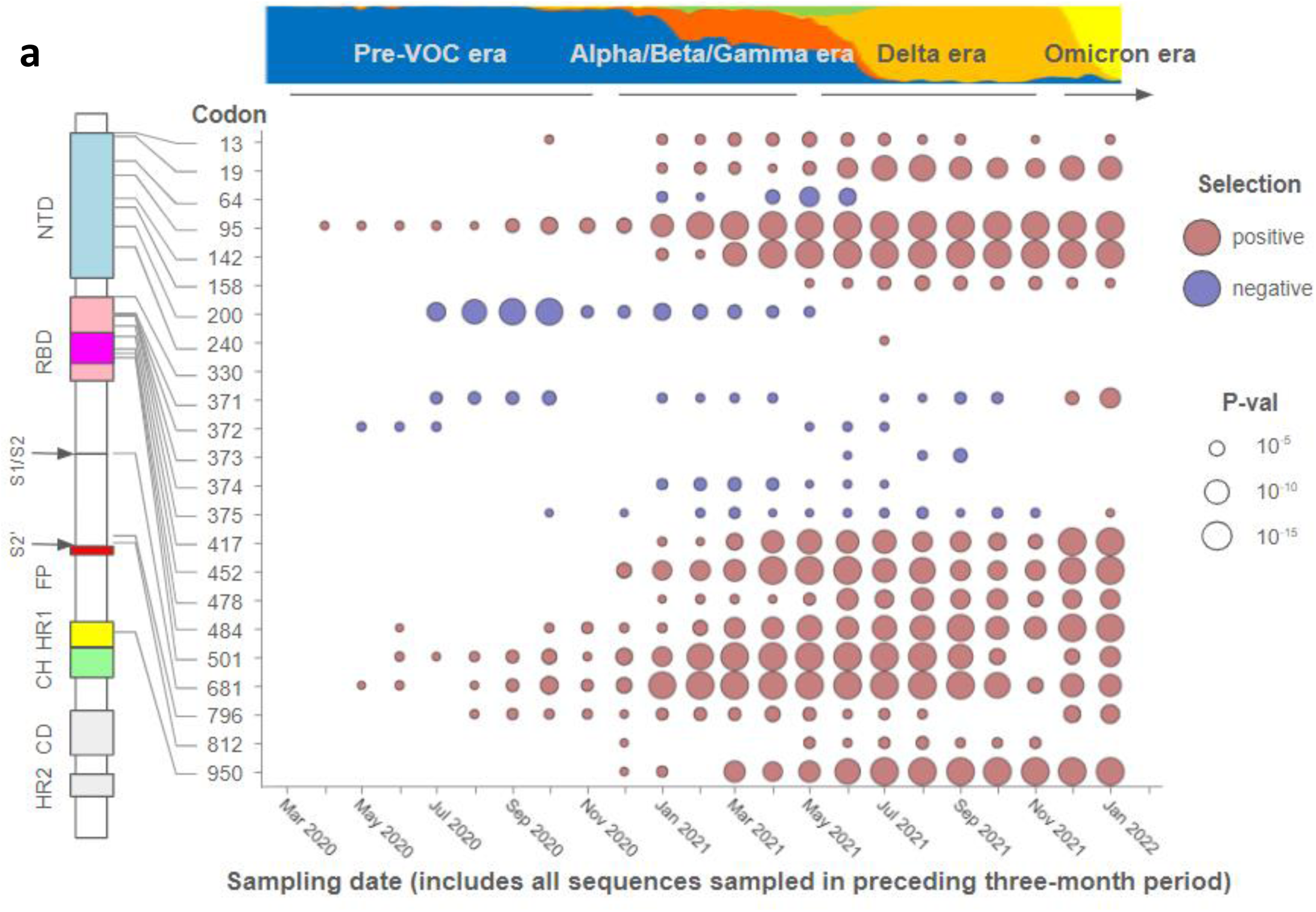
Signals of natural selection at a sample of SARS-CoV-2 S-Gene codon sites during the early phases of the COVID-19 pandemic. Evidence of positive (red circles) and negative (blue circles) was detected using the IFEL method using all globally sampled near-full length SARS-CoV-2 sequences in the three months preceding the indicated sampling dates with the sizes of circles indicating the degree of statistical support for the detected selection signals (see key on the right). The approximate era-during the COVID-19 pandemic (after Amodio et al) is indicated at the top. The non-variant of concern (VOC) lineages indicated in blue, the Alpha VOC in Red, the Beta VOC in dark orange, the Gamma VOC in green, the Delta VOC in orange and the Omicron VOC in yellow. The approximate location in the Spike protein of amino acids encoded by the various codon sites is indicated on the left: NTD = N-terminal domain; RBD = Receptor binding domain; FP = Fusion peptide; HR1 = Heptad repeat 1; CH = Central helix region, CD = Connector domain; HR2 = Heptad repeat 2.

**Supplementary Figure 5:**
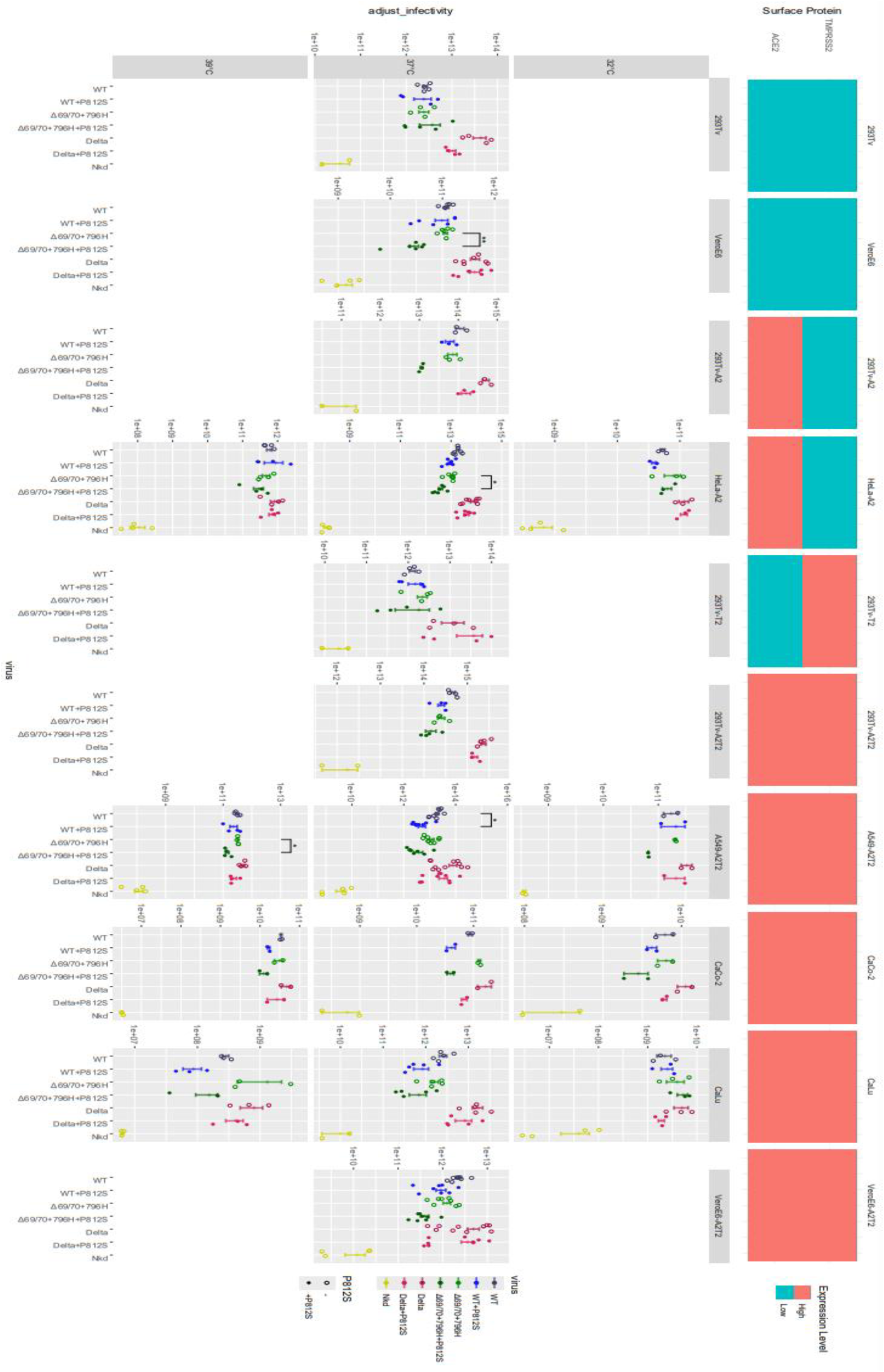
P812S has a detrimental effect on cell entry in additional cell lines across 32°C, 37°C, and 39°C. PV entry of WT (Wuhan-Hu-1+D614G), WT+P812S, WT+Δ69/70+796H, WT+Δ69/70+796H+P812S, Delta, Delta+P812S introduced to cells of varying levels of ACE2 and TMPRSS levels pre-incubated at 32, 37, and 39 degrees. Data are supplemented by mean ± standard error of n = 3–8 replicates. Statistical analysis was performed using unpaired student’s t-test with Benjamini-Hochberg correction.

**Supplementary Figure 6.**
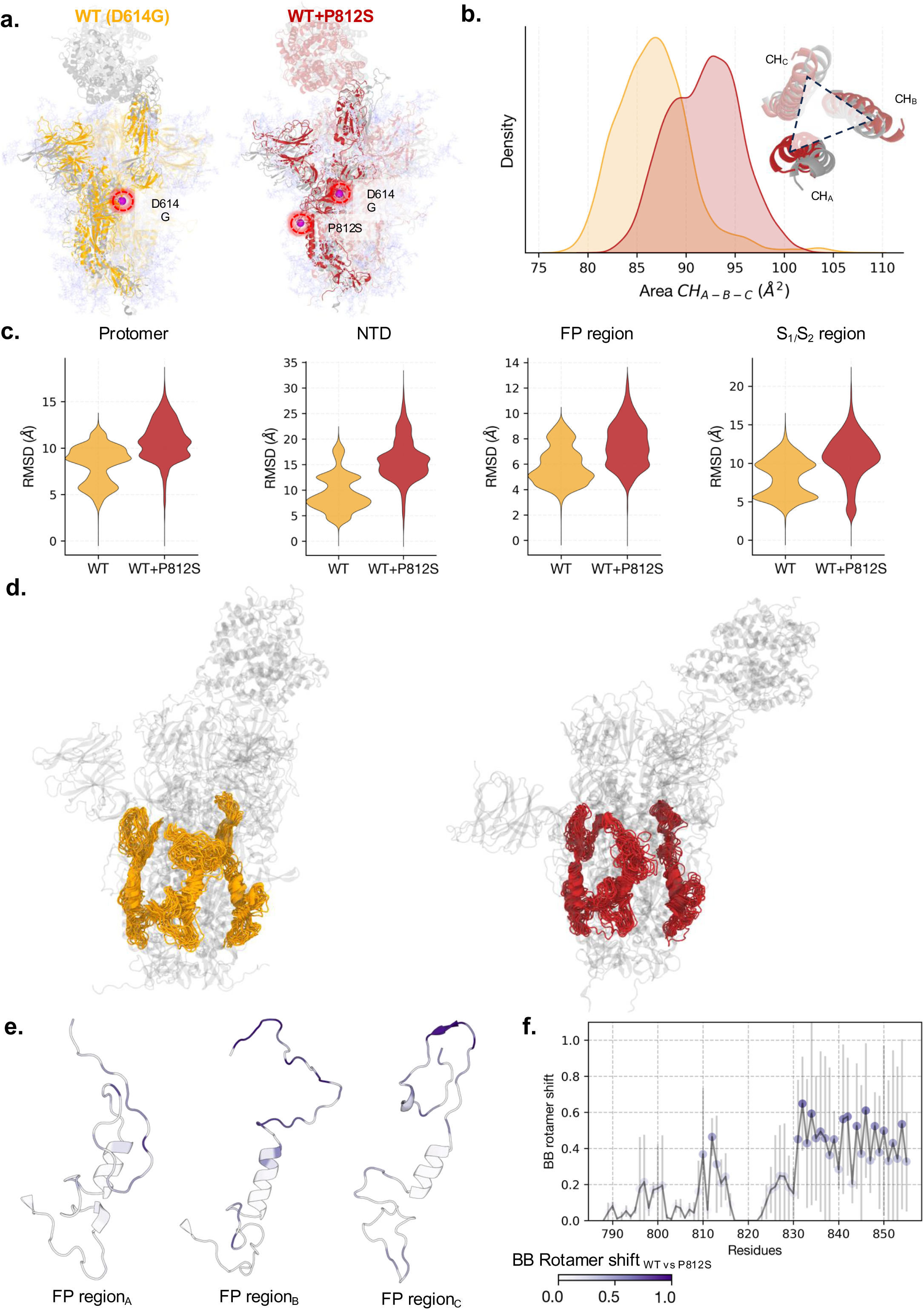
The P812S mutant impacts the dynamics at both the S1/S2 and FP/S2’ cleavage sites of Spike protein. a) Snapshot of WT (D614G) and P812S spike trimers in complex with ACE-2. Both 0ns (grey) and 1μs timepoints superimposed on each other. b) The probability density plot of the triangle area formed by the centers of three CH domains is compared between WT and P812S mutant. The snapshot highlights the eagle-eye view of inter-CH orientation in WT (grey) and P812S mutant (dark red). c) The violin plots compare the distribution of RMSD calculated for the entire protomer, NTD, FP and S1/S2 regions respectively. D) The snapshots highlight the ensemble of FP conformations in WT (left) and P812S mutant (right). e) The backbone rotamer shift calculated between WT and P812S is mapped on the individual FP regions of the spike trimer. f) The co **r**esponding rotamer shift values are plotted for FP residues. The values are averaged across three chains.

**Supplementary Figure 7:**
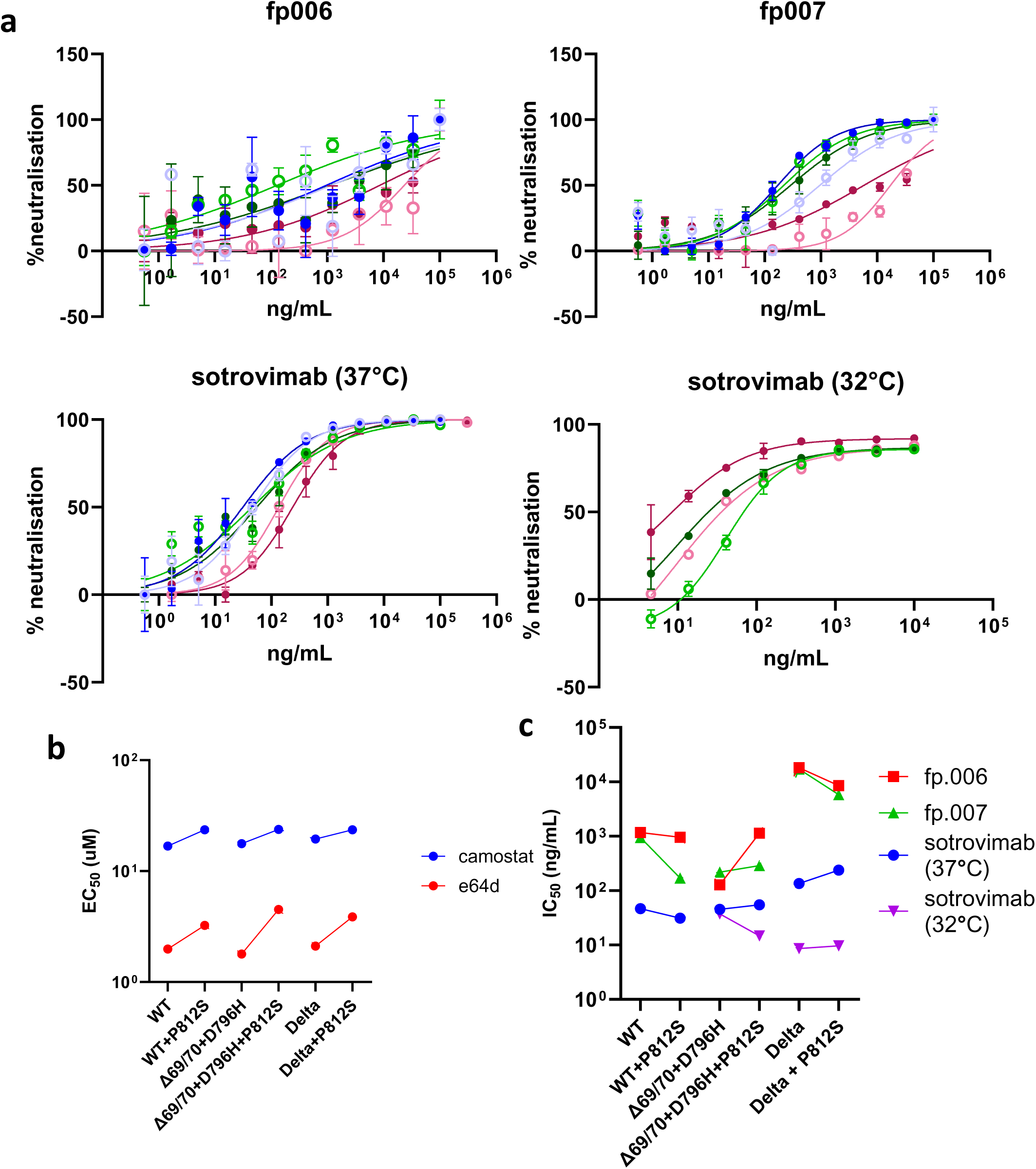
Individual curves of monoclonal antibodies. **a)** Individual inhibitor curves of monoclonal antibodies. b) EC50 of E64d (cathepsin inhibitor, blue) and camostat (TMPRSS2 inhibitor, red) in A549-ACE2/TMPRSS2 using PV expression respective Spike. c) IC50 of sotrovimab incubated with pseudovirus at 32°C (purple), sotrovimab at 37°C (blue), fp.006 (red), and fp.007 (green) neutralizing pseudoviruses corresponding to the indicated mutations. Results are representative of 2 technical replicates Data points are representative of 2 technical replicates. Error bars represent SD.

